# Vasopressin acts as a synapse organizer in limbic regions by boosting PSD95 and GluA1 expression

**DOI:** 10.1101/2020.11.07.373027

**Authors:** Limei Zhang, Teresa Padilla-Flores, Vito S. Hernández, Mario A. Zetter, Elba Campos-Lira, Laura I. Escobar, Robert P. Millar, Lee E. Eiden

**Author notes:** Corresponding authors: Limei Zhang, Department of Physiology, School of Medicine, National Autonomous University of Mexico, Av. Universidad 3000, CDMX, 04510, Mexico; Lee E. Eiden, Section on Molecular Neuroscience, NIMH-IRP, NIH, Bethesda, MD, USA. These authors contributed equally to this work (Limei Zhang, Teresa Padilla-Flores, Vito S. Hernández, Mario A. Zetter).

## Abstract

Hypothalamic arginine vasopressin (AVP)-containing magnocellular neurosecretory neurons (AVPMNN) emit collaterals to synaptically innervate limbic regions influencing learning, motivational behaviour, and fear responses. Here, we characterize the dynamics of expression changes of two key determinants for synaptic strength, the postsynaptic density (PSD) proteins AMPAR subunit GluA1 and PSD scaffolding protein 95 (PSD95), in response to *in vivo* manipulations of AVPMNN neuronal activation state, or exposure to exogenous AVP *ex vivo*. Both long term water deprivation *in vivo*, which powerfully upregulates AVPMNN metabolic activity, and exogenous AVP application *ex vivo*, in brain slices, significantly increased GluA1 and PSD95 expression measured by western blot, in brain regions reportedly receiving direct ascending innervations from AVPMNN (i.e., ventral hippocampus, amygdala and lateral habenula). In contrast, the visual cortex, a region not observed to receive AVPMNN projections, showed no such changes. Ex vivo application of V1a and V1b antagonists to ventral hippocampal slices ablated the AVP stimulated increase in postsynaptic protein expression measured by western blot. Using a modified expansion microscopy technique, we were able to quantitatively assess the significant augmentation of PSD95 and GLUA1 densities in subcellular compartments in *locus coeruleus*’ tyrosine hydroxylase immunopositive fibres, adjacent to AVP axon terminals. Our data strongly suggest that the AVPMNN ascending system plays a role in the regulation of the excitability of targeted neuronal circuits through upregulation of key post-synaptic density proteins corresponding to excitatory synapse.

**Supported by grants:** UNAM-DGAPA-PAPIIT-IN200121 & CONACYT-CB-238744 (LZ); CONACYT A1-S-8731 (LIE); MH002386, NIMH, NIH, USA (LEE)

## 1. Introduction

The neurohormone arginine vasopressin (AVP, also called antidiuretic hormone, ADH) is mainly synthesized in the hypothalamic paraventricular (PVN) and supraoptic (SON) nuclei by a type of cell with large somata (diameters around 20–35 μm) that are traditionally referred to as magnocellular neurosecretory neurons (AVPMNN) and is released mainly in the posterior hypophysis as the key regulator of body water homeostasis ^(1)^. The AVPMNN, together with those for oxytocin, were the first peptidergic systems to be described in the mammalian brain^(2)^. Release of AVP from both posterior pituitary lobe and median eminence, as well as AVPMNN somato-dendritic release within the hypothalamus itself ^(3, 4)^ helps to control water homeostasis.

David de Wied in the 1970s pioneered studies suggesting that vasopressin may act as a neurotransmitter or neuromodulator at synapses within the brain cognitive/emotional control centres ^(5, 6)^. Shortly after, a series of studies published in the 1980s and early 1990s showed that sub-micromolar concentrations of vasopressin could, in brain slices *ex vivo*, powerfully and reversibly increase the rate of firing of neurons in the CA1 areas of rat hippocampal slices and that this effect could be fully antagonized by an anti-vasopressor vasopressin analogue ^(7)^. Subsequently, AVP was localized to Gray type I synapses, usually corresponding to glutamatergic (excitatory) neurotransmission ^(8)^, in hippocampus ^(9)^, amygdala ^(10)^, lateral habenula ^(11)^ and locus coeruleus ^(12)^. These observations cemented the notion that AVP could act within the brain as well as in the periphery through AVP release from the same neurons, which therefore integrated regulation of homeostatic (hydromineral balance) and allostatic (behavioural) physiology within a single type of neurohormonal/neuropeptide cell ^(13, 14)^.

Despite the plethora of observations detailing AVP actions at hippocampal and other cognitive hubs in the brain (*vide supra*), the cellular mechanisms by which AVP potentiates neuronal excitability are not known. AVP-containing dense core vesicles (AVP+ DCV) have been observed at excitatory synaptic active zones, docked onto the presynaptic membrane, suggesting that vasopressin modulation of neurotransmission may occur at the level of the synapse itself within limbic regions such as hippocampus ^(9)^, amygdala ^(10)^, lateral habenula ^(11)^ and locus coeruleus ^(12)^. Figure 1, summarizes previous works, showing this phenomenon in lateral habenula, A, and ventral hippocampus, B, as two examples. In both cases, the animals were under food and water *ad libitum* intake condition before fixation.

**Figure 1.**
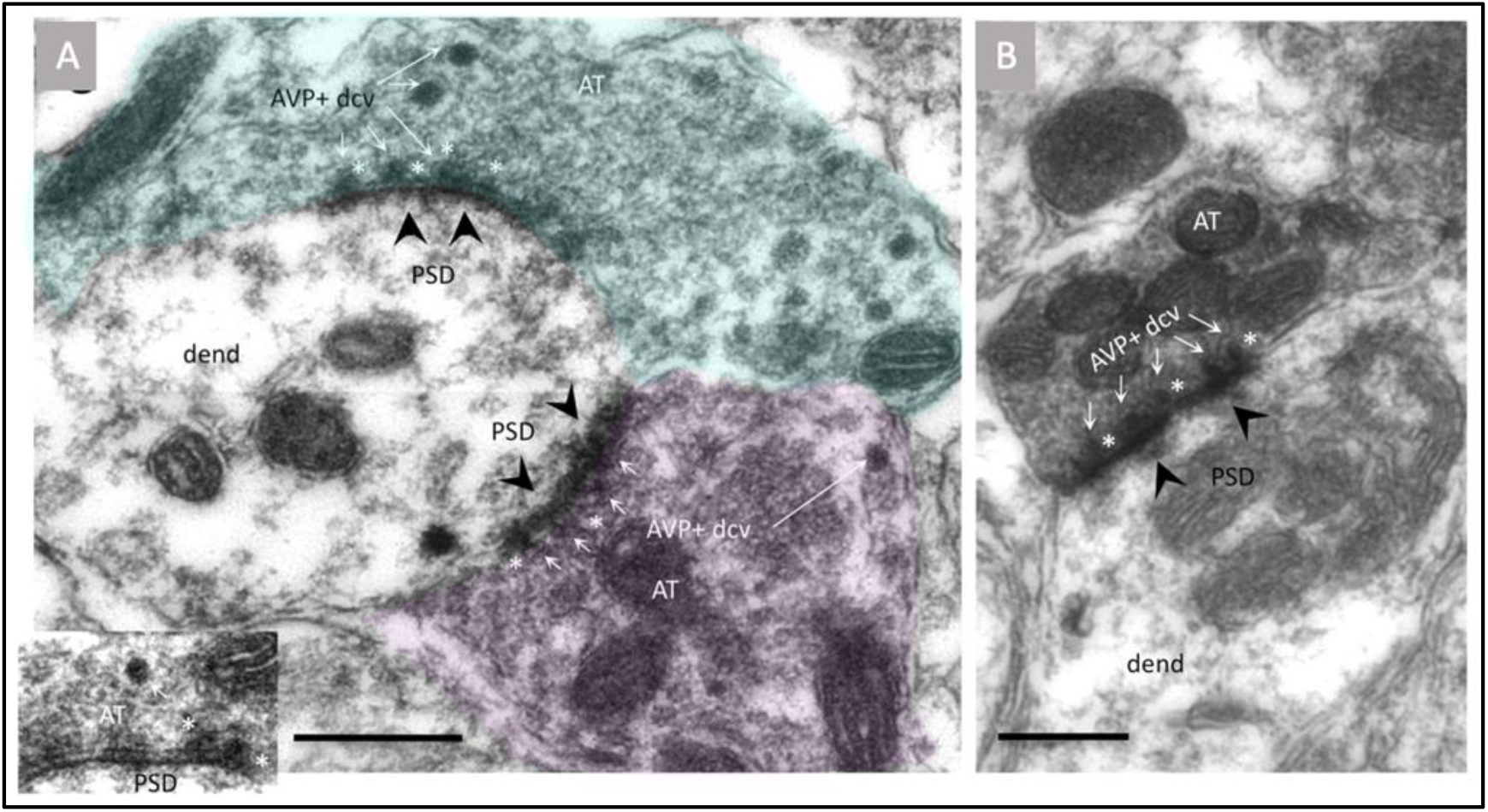
Ultrastructural examples showing morphological evidence that AVP could directly modulate synaptic function. **A:** two Gray type I synapses made by two AVP-immunopositive (AVP+) axon terminals (AT, green and pink) onto an unlabelled dendrite (dend) contain AVP+ dense core vesicles (DCV, white arrows). Inset shows higher magnification of four AVP+ DCV docked on the active zone of the presynaptic membrane. Postsynaptic density (PSD), a morphological feature of glutamatergic synapse, is indicated by black arrowhead. Sample was taken from rat lateral habenula. **B**: analogue notations to A, sample taken from rat ventral hippocampus. In both cases, the animals were under food and water *ad libitum* intake conditions before fixation. Panel A is modified from ^(11)^ (Creative Common Attribution License holder) and panel B is taken from ^(9)^ with permission from the original publisher. Scale bars: 500 nm.

With this background, we asked whether AVP could participate in the modulation of synapse strength via dynamic modification of the protein composition of the postsynaptic density (PSD). We characterized the dynamics expression changes of two PSD proteins, AMPAR subunit GluA1 and PSD scaffolding protein 95 (PSD95), which are known to be key determinants for synaptic strength, under in vivo and ex vivo conditions. Both long term water deprivation *in vivo* which powerfully upregulates AVPMNN metabolic activity, and exogenous AVP application in *ex vivo* brain slices, significantly increased GluA1 and PSD95 expressions measured by western blot, in brain regions that receive direct ascending innervations from AVPMNN (i.e., ventral hippocampus, amygdala and lateral habenula, *vide supra*); whereas the visual cortex, a region not observed to receive AVPMNN projections, showed no significant differences. *Ex vivo* application of V1a and V1b antagonists to ventral hippocampal slices ablated the AVP stimulated increase in postsynaptic protein expression measure by western blot.

Efforts to localize the PSD95 and GluA1 immunopositive puncta in the post-synaptic segment, in an identified cell type, adjacent to AVP immunopeptide axons were further made. Due to the tight association of the axonal bouton and the dendritic spine which limits the clear distinction under conventional confocal microscopy, we took advantage of newly developed expansion microscopy (ExM) technique ^(15-17)^, which utilizes isotropic tissue swelling to expand the sample, so that the resolving power to visualize the PSD proteins under regular confocal imaging can achieve the necessary resolution required to quantitatively assess the PSD95 and GluA1 immunopositive puncta’ densities. We observed significant augmentation of PSD95 and GLUA1 densities in locus coeruleus, a region demonstrated to receive direct AVPMNN innervation ^(12)^, under water deprivation conditions, within an identified cell type (TH+ neurons) adjacent to AVP axon terminals.

## 2. Materials and methods

### 2.1. Rats

A total of 56 male Wistar rats of 250 +/-50 g were used in this study. Rats were obtained from the local animal vivarium and were housed three per cage under controlled temperature and illumination (12h/12h) with water and chow *ad libitum* unless otherwise indicated. All animal procedures were approved by the local research ethics supervision commission (license number CIEFM-079-2020).

### 2.2 Chemical and primary antibodies used in this study

Arginine vasopressin (MK I-23) for western blot experiment was kindly donated by Professor Maurice Manning ^(18)^, AVP receptor antagonists, V1b (SSR149415) and V1a (SR49059) were provided by Axon Medchem (Groningen, The Netherlands). Primary antibodies used for western blot experiments were: mouse monoclonal anti-PSD95 (UC Davis, clone K28/74, 1 mg/mL, 1:1000, CA, USA), guinea pig polyclonal anti-GluR1 (Frontiers Institute, GluA1-GP-Af380, 200μg/mL, Japan, https://nittobo-nmd.co.jp/pdf/reagents/GluA1.pdf; 1:1000), mouse monoclonal anti-GAPDH (Glyceraldehyde-3-Phosphate Dehydrogenase) (GTX627408, 1:2000, GeneTex, CA, USA); for immunohistochemistry: mouse monoclonal anti-PSD95 (UC Davis, clone K28/74, 1 mg/mL, 1:1000, CA, USA), guinea pig polyclonal anti-GluR1 (Frontiers Institute, GluA1-GP-Af380, 200μg/mL, 1:500, Japan, 1:1000), sheep anti-TH (Chemicon/Sigma, AB1542), rabbit anti-AVP ^(19)^.

Chemicals used for western blot (standard method) and for expansion microscopy (modified method) are listed in the corresponding segments of this section.

### 2.3 Experimental design

Three sets of experiments (noted as A, B and C) were performed for this study.

Experiment A used western blot method to evaluate, the changes in PSD95 and GluA1 protein expression levels in an *in vivo* water deprivation treatment model, in which the *variables* were different *time-points* of the *treatment*. With this physiological treatment, the hypothalamic vasopressin system is known to be upregulated in a relatively selective manner, even before a significant disruption of the body’s water homeostasis occurs ^(20)^.

Experiment B aimed to evaluate, through western blot, whether the *ex vivo*, pharmacological exposure of ventral hippocampal slices to AVP 100nM^(21)^, over different time courses, exerts similar effects on PSD95 and GLUA1 expression as in the WD study, and whether the exposure of *equimolar concentrations* of its specific antagonists for V1a and V1b receptors, could ablate the increase in expression induced by AVP.

Experiment C was designed to evaluate, through immunohistochemistry and expansion microscopy, whether water deprivation increases the PSD95 and GluA1 immuno-positive puncta’ densities in a given region innervated by AVPMNN direct projection. We chose pontine locus coeruleus, as an example of a region densely innervated by AVP-immunopositive fibres which further increases in immunolabeling density with water deprivation ^(12)^. The strong somatic/dendritic expression of tyrosine hydroxylase and the cell population homogeneity of LC, together with the dense AVP innervation, allowed us to successfully observe the post-synaptic cellular segments contacted by AVP immunopositive fibres, despite the immunofluorescent signals decay with expansion microscopy processing. It is worth mentioning that the optical resolution of conventional immunofluorescence confocal microscopy limits the quantification of postsynaptic density proteins. With the modified expansion microscopy (2-3 times), this quantification is achievable, while at the same time conserving the regional/cellular structure and ease of specimen handling for microscopical examination.

### 2.4 Water deprivation (WD) treatment and unit-subject definition for western blot assay

Twenty-four experimental subjects were divided in four time-points: control (water and food *ad libitum*), 24 hours of water deprivation (24h WD), 48 hours of water deprivation (48h WD) and 24 hours of water restoration after 48h WD (48h WD + 24h R).

To show the dynamic change of PSD95 and GluA1 as a *function of the treatment*, and to assure that the samples for each region possessed anatomical precision and sufficient protein amounts, we introduce the *unit-subject* concept that is defined as protein samples of *a given anatomical region* coming from *eight rats, distributed in four time-points*, which were assayed in a same western blot. The reported results in each time-point have *n=3 unit-subjects (six rats in total)*. The blots of the three unit-subjects, per each region are presented in its entirety in the Fig. 2, in order to allow the direct observation of the data, in addition of the numerical data and statistical analysis derived from it (statistic raw data and description can be found in supplementary information).

**Figure 2.**
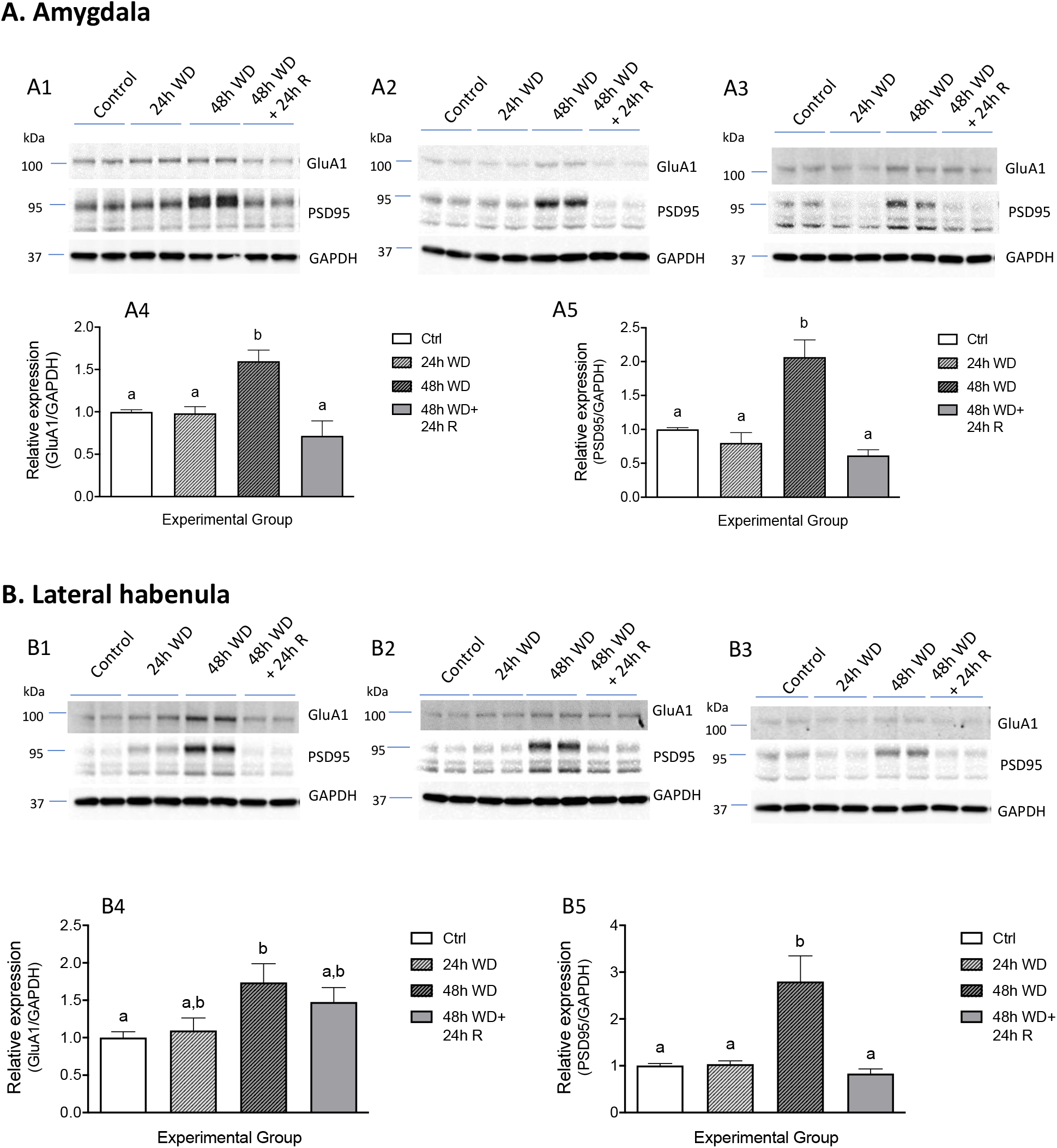
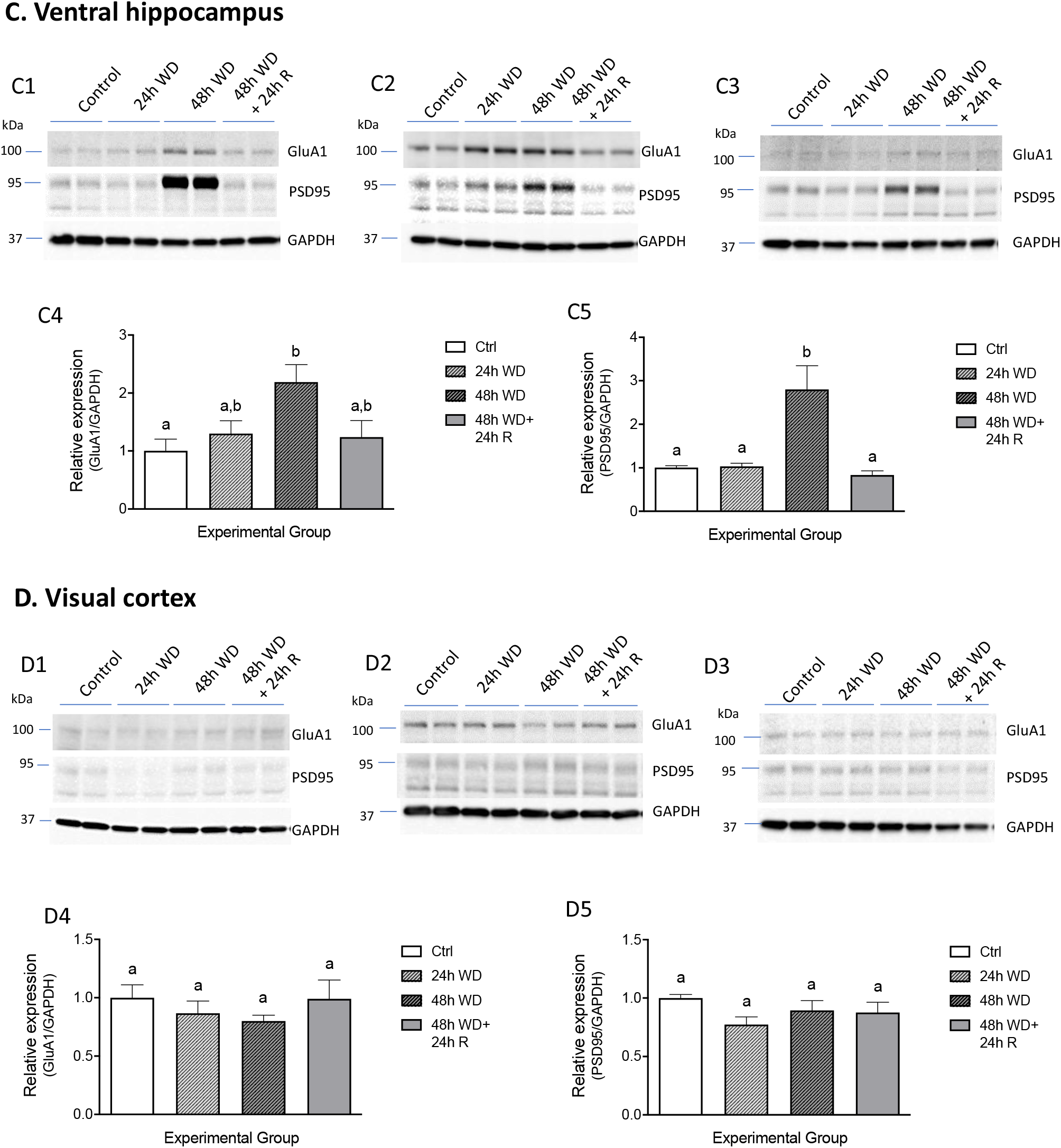
Prolonged water deprivation (48 hr WD) effectively increased GluA1 and PSD95 expression in the main AVPMNN’s ascending collaterals targeting regions: amygdala (A), lateral habenula (B); ventral hippocampus (C). We used the primary visual cortex (V1) as a negative control of the experiment (D). The mean relative expression for each treatment was normalized to the expression of the control levels. To show the individual variability and the consistence, as well as the direct basis of the statistic differences, we chose here to show the blots for each of the three unit-subjects (see methods section 2.5), each one includes the complete 4 time-point in duplicates. Bars that do not share a common letter, indicate significant differences between means.

### 2.5 Sample obtention and western blot procedure for in vivo experiments

After the corresponding treatment periods (control, 24h WD, 48h WD and 48h WD + 24h R), rats were deeply anesthetized with an overdose of sodium pentobarbital (180 mg/kg, Sedalpharma, México) and decapitated using a rodent guillotine. Fresh brain tissue was removed quickly and dissected on ice-cooled saline (0.9% NaCl) solution. Brains were coronally blocked from optic chiasm (around Bregma 0.00 mm) to posterior Bregma -7.00 mm and put on a tissue chopper for thick slice preparation (0.5 mm thickness). Segments of brain sections containing central amygdala, habenula, ventral hippocampus (vHi) and primary visual cortex (V1, served as a negative control region since no direct innervations to this region from hypothalamic AVPMNN system have been reported), were obtained under a stereoscopic microscope using microdissection tools. Each sample of a given brain region of interest in this study was pooled from the same region from two rats, to provide sufficient brain tissue for immunoblotting without sacrificing anatomical precision.

Dissected brain regions were sonicated in ice-cold extraction buffer: 50 mM Tris-HCl pH 7.4, 8.5 % sucrose, 2 mM EDTA, supplemented with proteases and phosphatases inhibitors (8 μg/mL aprotinin, 10 μg/mL leupeptin, 4 μg/mL pepstatin A, 500 μM AEBSF, 5 mM benzamidine, 20 mM ß-glycerophosphate, 10 mM sodium fluoride, 1 mM sodium orthovanadate). Brain homogenates were assayed for protein using the BCA method (Pierce, Thermo Scientific cat# 23225). For SDS-PAGE (sodium dodecyl sulfate polyacrylamide gel electrophoresis), homogenates (15 μg total protein) were mixed with loading buffer added with ß-mercaptoethanol and boiled for 3 min. Samples were separated by electrophoresis in 10% acrylamide gels and transferred to 0.2 μm nitrocellulose membrane (Amersham, GE cat# 10600006). Blots were blocked for 1 h at room temperature (RT) in Tris-buffered saline with 0.05 % Tween-20 (TBST) containing 5 % blotting-grade dry milk (Biorad cat# 170-6404), incubated overnight with primary antibody (dilutions above mentioned) in blocking buffer at 4°C, washed 3x 10 min in TBST, incubated with secondary antibody conjugated to horseradish peroxidase (1:5000, Jackson ImmunoResearch) for 1 hr at room temperature, and washed 3x 10 min in TBST. Membranes were visualized with luminol substrate by enhanced chemiluminescence ^(22)^. PSD95 and GluA1 were evaluated in the same western blots, for which membranes were incubated in stripping buffer (0.2 M glycine pH 2.2, 0.1% SDS, 1% Triton X-100) for 2x 15 min at room temperature, washed and re-probed using the above protocol. Antibodies detecting GAPDH or actin were used to provide loading controls

Blots were scanned with ChemiDoc (BioRad, USA). The images were converted to grayscale and the densitometric analysis was performed using FIJI Gel analizer tool (Image J, NIH, Bethesda, USA). Briefly, a rectangular selection was drawn around the bands detecting the postsynaptic proteins GluA1 or PSD-95 and a similar selection around the GAPDH band. In the case of PSD-95 two bands were observed, one at the expected molecular weight of 95 kDa and other around 70 kDa (https://neuromab.ucdavis.edu/datasheet/K28_74.pdf). Since the origin of this lower molecular weight band is controversial, we chose to only measure the band around 95 kDa. After selecting similar size regions, the program plots a graph representing the integrated optical density along the length of the selected area, the area under the curve represents the optical density of that band. The obtained optical density of PSD95 or GluA1 band was divided over the optical density of the GAPDH band, and the result of each condition was normalized with respect to the average value of the control bands (set to be 1), the results are therefore presented as relative expression of GluA1/loading control or PSD95/loading control.

### 2.6 Hippocampal acute slice pharmacology with exogenous AVP

Rats of approximately 200g were deeply anesthetized using an overdose of sodium pentobarbital (180 mg/kg, Sedalpharma, México) and killed by rapid decapitation using a rodent guillotine. Brain horizontal slices of 300 μm thickness containing the ventral hippocampus were obtained using a Leica VT 1000S vibratome, in ice-cold cutting solution containing the following (in mM): 75 NaCl, 2.5 KCl, 25 NaHCO_3_, 1.25 NaH_2_PO_4_, 25 glucose, 0.1 CaCl_2_, 6 MgCl_2,_ and 50 sucrose bubbled with a mixture of 5% CO_2_ and 95% O_2_, pH 7.4 ^(11)^. The slices between interaural coordinates 1.50 and 2.10 mm were selected and transferred to artificial cerebrospinal fluid (ACSF) containing the following (in mM): 125 NaCl, 2.5 KCl, 25 NaHCO_3_, 1.25 NaH_2_PO_4_, 25 glucose, 2 CaCl_2_, and 2 MgCl_2_, bubbled with a mixture of 5% CO_2_ and 95% O_2_, pH 7.4, and left to stabilize at room temperature for 30 min ^(11)^. After this time, the slices were incubated at 37ºC and added with the corresponding reagents in each experiment.

To determine the AVP’s role in the upregulated expression of GluA1 and PSD95, observed *in vivo*, we evaluated the *in vitro* effects of exogenous arginine vasopressin AVP applied to the slices in the incubation medium. Six rats were used in this experiment. In order to obtain sufficient amount of tissue for each blot, tissue from two rats was pooled: one hemisphere for control and one for AVP 100nM incubation. To determine the time course of exogenous AVP (100nM) application in the changes observed in the previous experiment, we assayed PSD95 and GluA1 expression under exogenous AVP application (ACSF alone, or ACSF with 100nM AVP at four time-points: baseline, 5 min, 60 min or 120 min. We used 8 rats for this experiment. Brain slices containing ventral hippocampus from the 16 hemispheres were distributed among 4 time-points and slices from two rats were pooled to run in the same blot. To assess if the effects were mediated by V1a and/or V1b receptors, we incubated slices from an additional 8 rats (16 hemispheres) during 60 minutes in ACSF alone, AVP 100nM, AVP 100nM plus 100nM SR49059 or AVP 100nM plus 100nM SSR149415. After the incubation, tissue was processed for western blot through SDS-PAGE electrophoresis as above mentioned.

### 2.7 Immunostaining and expansion microscopy (ExM)

Control and WD 48 hr male rats (n=5 and N=10) were euthanized as described and perfused transaortically with 0.9% saline followed by cold fixative containing 4% of paraformaldehyde in 0.1 M sodium phosphate buffer (PB, pH 7.4) plus 15% v/v saturated picric acid for 15 min. The brains were sectioned using a Leica VT 1000S vibratome, at 60μm thickness in the horizontal plane.

Sections containing locus coeruleus (LC) were blocked with normal donkey serum (NDS, 20%) in Tris buffered-saline (TBS, pH7.4) plus 0.3% Triton-X (TBST) for 1 hour and then incubated with the following primary antibodies: sheep anti-TH, rabbit anti-AVP, mouse anti-PSD95 and guinea pig anti-GluA1 diluted in TBST plus 1% NDS over two nights, followed by incubation with corresponding secondary fluorochrome-conjugated antibodies and mounted on glass slides with fluorescent mounting medium and coverslipped. For GluA1 antigen retrieval, slices were first treated with pepsin 1mg/ml diluted in 0.2M HCl, and incubated during 10 minutes at room temperature ^(23)^

After conventional imaging, the selected slices were unmounted and processed through a modified expansion microscopy (ExM) technique based on previous protocols ^(15-17)^. Briefly, after washing the slices with PBS (3×15 min at 4°C), slices were anchored with Acryloyl-X (1:100) in NaHCO3 (150mM) buffer overnight at room temperature. Sections were then washed with PBS and embedded in gelling solution (monomer solution^(15-17)^, 4-hydroxy-(2,2,6,6-tetramethylpiperidin-1-yl)oxidanyl(TEMPO), tetramethylethylenediamine (TEMED) and ammonium persulfate in proportion 47:1:1:1) for 30 minutes at 4°C. Sections were then immersed in gelling solution in a microscope slide with parafilm, covered with a coverslip and incubated at 37°C for two hours. Gel-embedded sections were trimmed to remove excess of gel and digested using 50 mM Tris buffer plus 0.8 M guanidinium chloride, 8 U/ml proteinase K, 0.5% Triton X-100, pH 8.0, for 3 hours at room temperature. Samples were then washed with PBS (0.28 M, 1 tablet of Sigma P4417 in 200ml of distilled water), no expansion was observed at this time point. When distilled water was used for subsequent washes (3 washes, 20 min each), the sample expanded up to 6 times. This ratio of expansion is not adequate for our experimental aims since the target region and signals are largely dispersed making imaging process difficult. Thus, a modified solution of PBS 0.14M (half of the concentration of the conventional PBS) was used, obtaining an expansion of approximately 3 times, which made possible the numerical counting of the PSD95 and GluA1 immunopositive puncta while preserving the neural structure. The samples were flattened in a coverslip treated with poly-L-lysine, with the face of the gel to be imaged toward the coverslip, then mounted in a constructed chamber filled with aqueous mounting medium with the same osmolarity to avoid shrinking.

### 2.8 Imaging and quantification of PSD95 and GluA1

Samples were examined in an inverted confocal microscope (Zeiss LSM 880, Germany). Images were taken with an optical section of 1 AU. To determine the expansion factor, nuclei of TH positive cells were taken as a reference and diameters were measured in 5 cells from pre-expanded and 5 cells from post-expanded samples.

For quantification of PSD95 and GluA1 immunopositive puncta, confocal images taken with 63X objective on expanded samples were assessed by 3 investigators blind to experimental conditions. Counting was made within an area of 100 μm^2^ (expanded area) and data were analysed for normality and differences obtained with unpaired *t*-test.

### 2.9 Statistical Analysis

Quantitative results were expressed as mean ± SEM. The Shapiro-Wilk test was used to confirm their normal distribution. To evaluate differences between means we used Student t-test or one-way ANOVA followed by Tukey post-hoc test. All the analyses were performed using Prism (GraphPad Software, V8.0, San Diego, CA, USA). The significance level was set at *p*<0.05. Raw data and results from all the statistical test performed are included in supplemental material. We used letters over the bar-graphs to depict significant differences between means. Groups with shared letter(s) indicate no-significant differences.

## 3. Results

### 3.1 . Water deprivation increased PSD95 and GluA1 protein expression in amygdala, lateral habenula and hippocampus

Water deprivation for 48 hours has previously been shown to activate the AVPMNN system ^(11)^. Western blot analysis revealed that 48 hours of water deprivation produced a significant increase in amount of protein expression of Glu1A and PSD95 in amygdala, lateral habenula, and ventral hippocampus, three regions that had previously demonstrated to receive direct innervation from the hypothalamic AVPMNN system ^(9-11, 24)^. In amygdala, three unit-subjects (see section 2.5) blots were displayed as A1-A3. The consistent increased protein levels of PSD95 and GluA1 were observed after 48 hr of water deprivation (Fig. 2, A1-A3). Bar graphs (panels A4 and A5) show the quantification of the relative protein expression levels compared to control. One-way ANOVA showed that treatment significantly affected the expression of GluA1 (F (3,20) = 10.09, *p*<0.001) and PSD95 (F (3, 20) = 17.70, *p*<0.001). Tukey multiple comparison test showed significantly higher levels of GluA1 after 48h WD (1.60 ± 0.129) compared to control (1.00 ± 0.025), 24h WD (0.983 ± 0.079), and 48h WD + 24 R (0.716 ± 0.177). PSD95 expression was also increased after 48h WD (2.067 ± 0.253) compared to control (1.00 ± 0.025), 24h WD (0.8 ± 0.152), or 48h+R (0.617 ± 0.083).

Likewise, in lateral habenula (Fig. 2, B panels), WD treatment significantly affected the expression of GluA1 (F (3,20) = 3.479, *p*<0.05) and PSD95 (F (3, 20) = 9.050, *p*<0.001). Tukey multiple comparison test showed increased levels of GluA1 expression after 48h WD (1.739 ± 0.250) compared to control (1.00 ± 0.079); PSD95 expression was increased after 48h WD (2.901 ± 0.524) compared to control (1.00 ± 0.033) to 24h WD (1.167 ± 0.244) or 48h+R (1.104 ± 0.170). In ventral hippocampus, (Fig. 2 panels C), correspondingly, 48 h WD treatment significantly affected the expression of GluA1 (F (3,20) = 4.154, *p*<0.05) and PSD95 (F (3,20) = 10.76, *p*<0.001). Tukey multiple comparison test showed significantly higher levels of GluA1 expression after 48h WD (2.188 ± 0.3) compared to control (1.00 ± 0.201); PSD95 expression was also increased after 48h WD (2.8 ± 0.549), compared to control (1.00 ± 0.044); 24h WD (1.033 ± 0.071) or 48h+R (0.833 ± 0.098). In contrast, in the visual cortex (Fig. 2, D panels), where AVPMNN vasopressin direct innervation has not been observed, no significant differences in either PSD95 or GluA1 expressions were observed (see supplemental materials for detailed statistical table and descriptive results). In all three regions we studied, a restoration of the expression of GluA1 and PSD95 to control levels was noted after 24 h recovery from WD.

To show both the individual variability, the consistency among the three *unit-subjects*, and the reproducibility of the conclusions drawn from the experimental observations, as well as their statistical differences, we chose to show the original western blot sets of all three *unit-subjects* in this report.

### 3.2 PSD95 and GluA1 protein expression in ventral hippocampus (vHi) slices were increased after incubation with AVP

To assess whether the AVPMNN vasopressinergic innervation to the brain limbic regions assessed above played a prominent role in upregulating the GluA1 and PSD95 expression levels after water deprivation, we performed an *ex vivo* pharmacological experiment in acute ventral hippocampal slices incubated during 1h in oxygenated ACSF solution with 100nM vasopressin.

We selected the ventral hippocampus due to the high level of AVP immunopositive fibres reported in a previous study in which the AVPMNN system was identified as one of the sources of their fibres ^(9)^. Each blot was constructed from the tissue provided by two rats (one hemisphere for each condition). After one hour of incubation with 100 nM AVP, we carried out western blot analysis of the expression of GluA1 or PSD95 protein levels. A two-tailed Student’s *t*-test showed that AVP treated slices increased their expression of GluA1 (control: 1.00 ± 0.045 *vs*. AVP: 1.67 ± 0.117, *p*<0.01) and PSD95 (control: 1.00 ± 0.068 *vs*. AVP: 1.83 ± 0.219, *p*<0.05). Three blots and quantification of the relative protein expressions with respect to control are depicted in Fig. 3 A.

**Figure 3.**
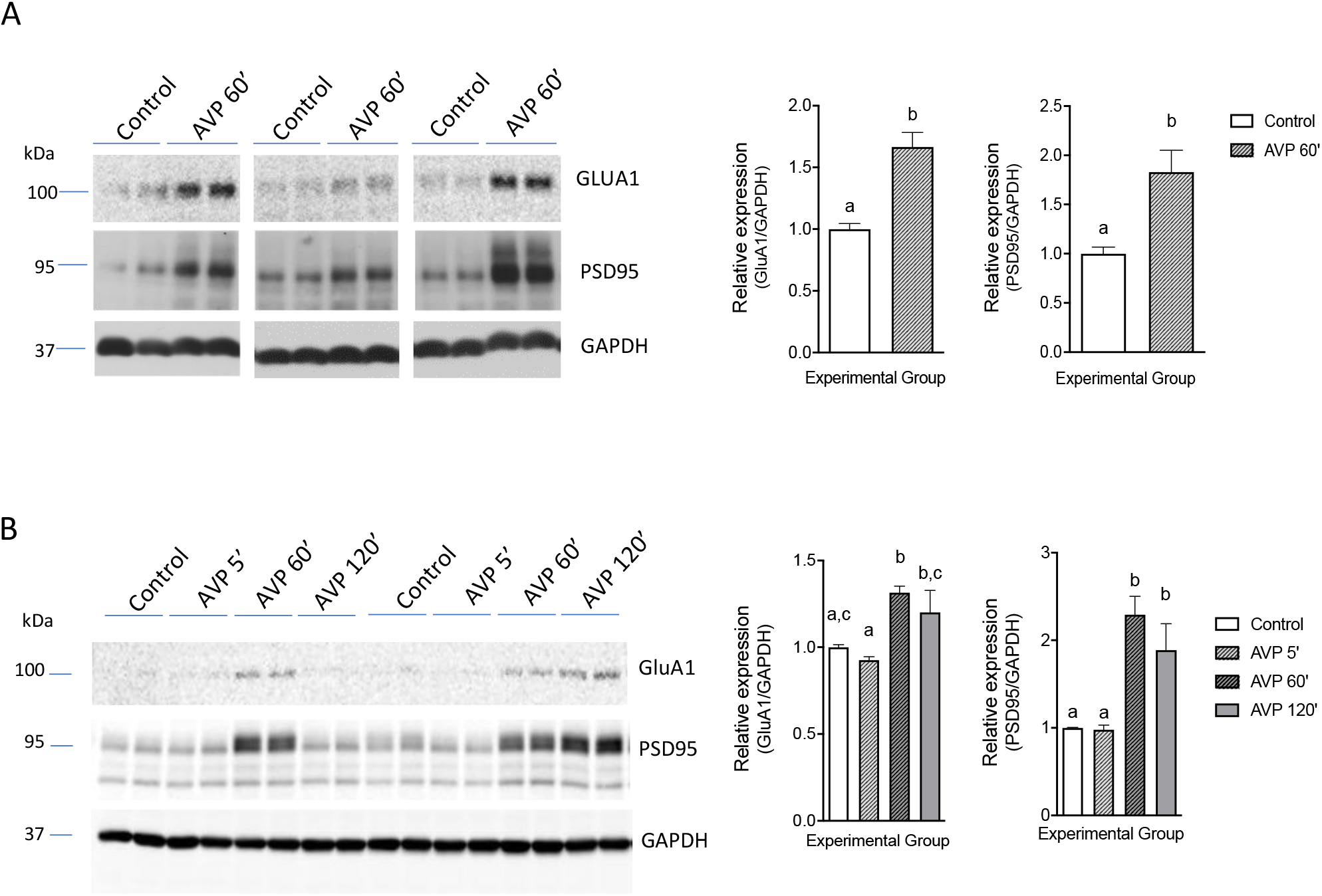
AVP increased the expression of GluA1 and PSD95 *ex vivo*. Panels A: GluA1 (*upper blot)*, PSD95 (*middle blot)* and GAPDH (*lower blot*) from lysates of ventral hippocampus formation dissected from acute brain slices (thick 300 μm) obtained from 6 rats (two rat samples per each condition), were incubated during two hours in oxygenated ACSF with vasopressin 100 nM. Panels B: show the relative expression of GluA1 and PSD95 in four times points (tissue from two rats in each lane). Common letter(s) indicate no-significant differences between means.

To assess the time course of the AVP effects on postsynaptic proteins expression, we compared ventral hippocampus slices incubated in oxygenated ACSF (control) or exposed to 5, 60 or 120 minutes of ACSF with addition of 100nM AVP. A total of eight rats provided the tissue for this experiment (four hemispheres per condition). Two-way ANOVA showed an effect of treatment for GluA1 (F (3,28) = 7.049, *p*<0.01) and for PSD95 (F (3,28) = 17.5, *p*<0.001). Post-hoc Tukey multiple comparison test showed a significant increase of GluA1 at 60 minutes of AVP exposure (1.315 ± 0.037) compared to control (1 ± 0.015). For PSD95, significantly higher expression levels were found after 60 min (2.29 ± 0.179) and 120 min of AVP exposure (1.89 ± 0.251) compared to control (1 ± 0.024). Quantification of the relative protein expressions for each time point with respect to control are depicted in Fig. 3B.

### 3.3 Vasopressin induced increase of GluA1 and PSD95 expression in ventral hippocampus (vHi) slices is blocked after incubation with AVP receptor antagonists

Since both V1a and V1b vasopressin receptors have been reported to be present in the hippocampus^(25, 26)^. We investigated whether antagonists for either receptor could revert the AVP-induced increase in GluA1 and PSD95 expression. Eight rats were used to obtain ventral hippocampus slices for incubation in ACSF (control), AVP 100nM, AVP 100nM + SR49059 100nM, or AVP 100nM + SSR149415 100nM. Following a 60min incubation period, GluA1 and PSD95 levels were assessed by western blot and their relative expression (normalised to control) was compared by a two-way ANOVA. Significant effect of treatment was observed for GluA1 (F (3,28) = 3.664, *p*<0.05) and PSD95 (F (3,28) = 28.19, *p*<0.001). Tukey’s Multiple comparisons test showed that compared to control, AVP treated slices had significantly increased expression of GluA1 (AVP: 1.652 ± 0.455 vs control: 1 ± 0.132) and PSD95 (AVP: 1.514 ± 0.046 vs control: 1 ± 0.161). Slices incubated with AVP and equimolar concentrations of V1a antagonist (SR49059), or V1b antagonist (SSR149415)) blocked the effects of AVP, although the effect on GluA1 was less prominent than that on PSD95 (Fig. 4).

**Figure 4.**
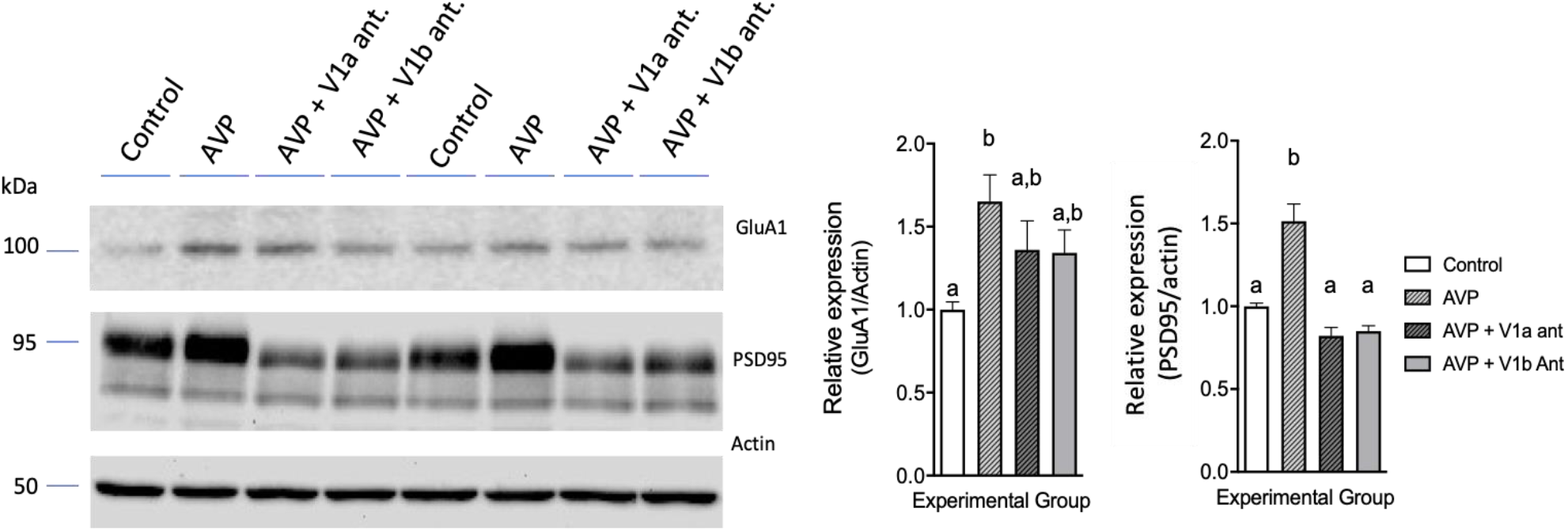
AVP induced increase in the expression of GluA1 and PSD95 *ex vivo* was blocked by V1a and V1b receptor antagonists. Left: GluA1 (*upper blot)*, PSD95 (*middle blot)* and actin (*lower blot*) from lysates of ventral hippocampus formation, dissected from acute brain slices obtained from 8 rats (16 hemispheres), which were incubated during one hour in oxygenated ACSF with vasopressin 100nM (AVP), AVP plus the V1a receptor selective antagonist (SR49059) or AVP plus the selective V1b receptor antagonist (SSR149415). Left bar graph: relative expression (mean ± SEM) of GluA1. Right bar graph: relative expression (mean ± SEM) of PSD95. Bars that do not share a letter indicate significant differences between means.

### 3.4 WD increased the PSD95 and GluA1 immunopositive puncta’s densities in TH+ dendritic compartment in locus coeruleus: puncta in apposition to AVP+ axonal varicosities

After observing the augmentation in the amounts of PSD95 and GluA1 following water deprivation *in vivo* and AVP *ex vivo* exposure, with western blots, we sought to evaluate whether water deprivation increases the PSD95 and GluA1 immuno-positive puncta’s densities in a given brain region innervated by AVPMNN direct projections. We chose pontine *locus coeruleus* (LC), as an example of a region densely innervated by AVP-immunopositive fibres which increases with water deprivation (Fig. 5, A and B, and also see the reference ^(12)^). The rationale of this selection is that the strong somatic/dendritic expression of tyrosine hydroxylase (TH) of the noradrenergic neurons in LC and its cellular homogeneity would help us to overcome the difficulties caused by immunofluorescent signal decay during the ExM processing. It is worth mentioning that the optical resolution of conventional immunofluorescence confocal microscopy limits the clear quantification of postsynaptic density proteins. Figure 5C shows a TH immunopositive neuron (diameter of nucleus: 7μm) and its inset showing the triple immunofluorescence reaction, before ExM processing, where the signals from TH, AVP and GluA1, blue, red, and green, respectively, are (partially, at least) overlapping, indicated with pink arrows. With the modified expansion microscopy (2-3 times, see method section 2.7), this quantification is achievable, whereas conserving the regional/cellular structure and easy handling for microscopical examination. Figure 5D shows an expanded TH+ positive neuron (diameter of nucleus: 21μm) with well conserved somato-dendritic cytoarchitecture. Inset, taking from the same sample, shows clear separation of the three immunofluorescence signals (TH, AVP and GluA1, blue, red and green, respectively, indicated with corresponding arrows). Appositions between AVP axons (red), TH+ dendrites containing GluA1 / PSD95 puncta (green) are depicted in Fig. 5 E -G F respectively. Panels E and E’ are photomicrographs of TH/AVP/GluA1 or TH/AVP/PSD95 respectively, from control rats. Panels F1-F4 and G1-G4) are taken from the same regions of F and G, of rats that underwent 48 hr of water deprivation, with further digitally magnified, in different Z levels, showing the abundance of these two PSD proteins in dendritic compartments. AVP axonal varicosity in apposition to TH+ dendrite, containing PSD proteins are indicated with asterisks. This latter aspect is exemplified at ultrastructural level with Fig. 5H, which is a representative electron micrograph taken from reference ^(12)^, showing AVP containing axon terminal establishing Gray type I synapse on the TH+ dendrite. A comparison of GluA1 and PSD95 puncta’s density in locus coeruleus TH+ dendritic compartments was performed on expanded immunoreacted slices from rats (n=5, N=10) that underwent 48h of water deprivation (WD 48hr) against those that were under food and water ad libitum conditions. Student *t-test* showed that WD48hr produced a significant increase in the density of GluA1 (WD 48hr: 10.7 ± 0.633 *vs* 6.8 ± 0.633, *p*<0.001) and PSD95 (WD 48hr: 16.5 ± 0.992 *vs* 9.9 ± 1.22, *p*<0.001) puncta per 100μm^2^ of TH+ dendritic compartments on the expanded confocal (1 Airy unit = 0.68 μm optical section thickness) photomicrograph. Results of the comparison are shown as mean ± SEM in the bar graph of Fig. 5I and J.

**Fig 5:**
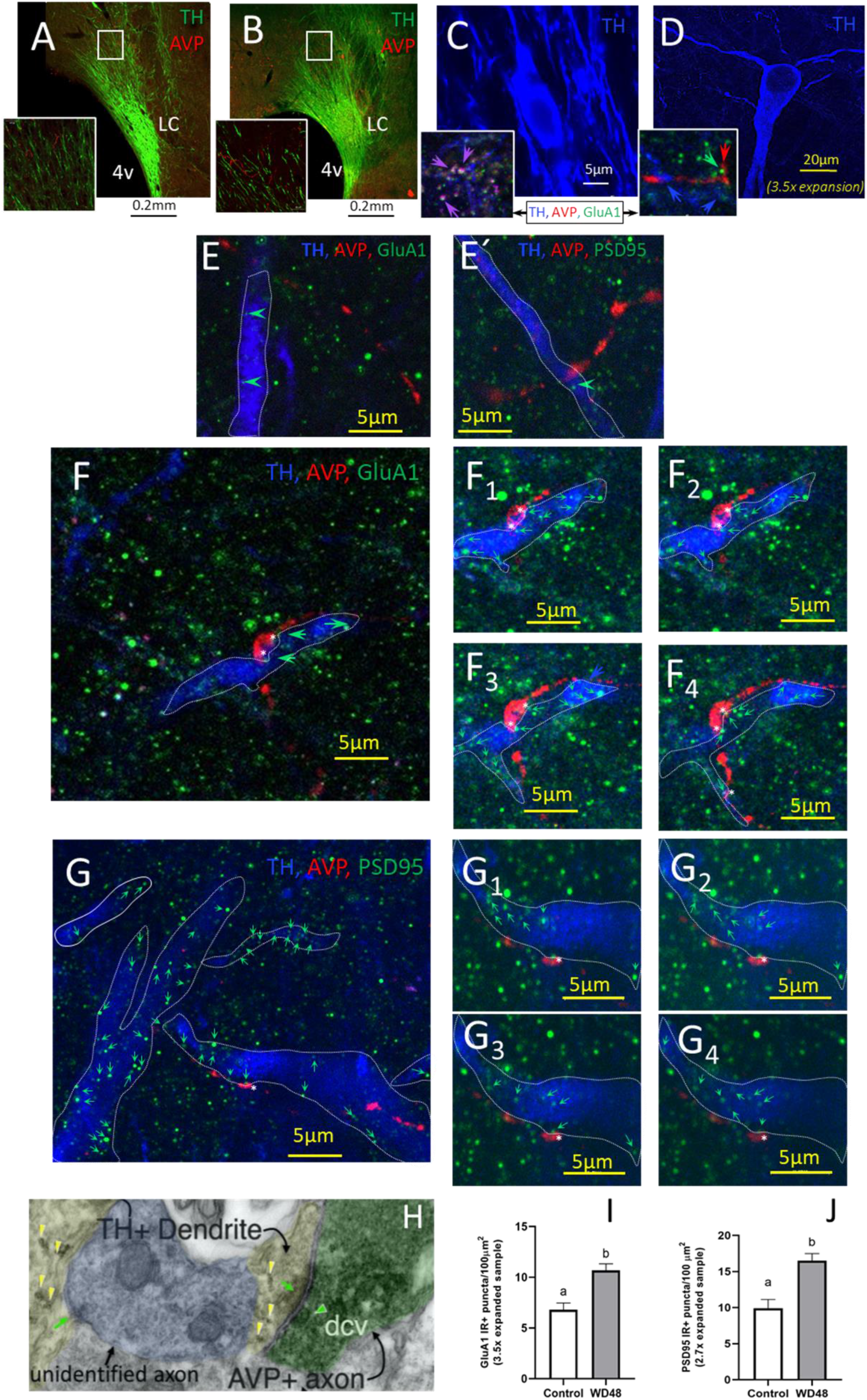
Water deprivation increased the density of GluA1 and PSD95 immunopositive puncta in the tyrosine hydroxylase immunopositive dendritic compartment in locus coeruleus. Panels A and B show representative locus coeruleus photomicrographs in control and 48WD rats. Water deprivation for 48 hours (WD48) increased the AVP immunoreactivity (red) located in the region where dendrites immunopositive for tyrosine hydroxylase (TH, green) are abundant. C and D: confocal photomicrographs (63X objective was used in both case) of TH immunopositive neurons before (C) and after (D) expansion microscopy (ExM). An increase of approximately 3 times the soma size (N.B: diameters of the nucleus increased from 7 μm to 21 μm) was observed after expansion (white or yellow scale bars indicate pre-ExM and post-ExM photomicrographs, respectively. In the high magnification confocal (1 Airy unit) images depicted in the insets, that before ExM (C), the GluA1 puncta are confluent (pink arrows) and after ExM (D) the GluA1 puncta (green) are separated from other two-coloured signals (red, green, and blue arrows indicate AVP, GluA1 and TH immunopositive compartments, respectively) N.B.: the background noise is also lowered by ExM processing. Panels E - G are examples of post-ExM photomicrographs of GluA1 (Es) or PSD95 (Fs) immunoreacted sections respectively. Panels E and E’ are photomicrographs of TH/AVP/GluA1 or TH/AVP/PSD95 respectively, from control rats. Sub-panels (F1-F4 and G1-G4) are taken from the same regions of F and G, with further digital magnification, in different Z levels, showing the abundance of these two PSD proteins in TH+ dendritic compartments. AVP axonal varicosity in apposition to TH+ dendrite, containing PSD proteins are indicated with white asterisks. H: electron photomicrograph shows ultrastructural evidence of the existence of putative glutamatergic (asymmetric) synapses between AVP+ axon and TH immunopositive dendritic segment within the locus coeruleus. Green arrowheads show dense core vesicles typical of peptidergic transmission (taken from reference ^(12)^, Creative Common Attribution License holder). I-J: Bar graphs show the means ± SEMs of the GluA1 and PSD95 puncta density in control and WD48 rats, respectively. Quantification was made after ExM within TH+ dendritic compartment in the *locus coeruleus*. Different letters above the bars indicate significant differences between groups. Scale bars for expanded samples are yellow.

## 4. Discussion

In this study, we investigated the dynamic changes of PSD95 and GluA1 expression levels and their localization in response to WD and AVP. Our data unambiguously demonstrated the augmentation of these post-synaptic density (PSD) proteins amounts in 48 hours water deprived animals, in brain limbic regions receiving direct ascending projections from the hypothalamic vasopressinergic magnocellular neurosecretory neurons (AVPMNN). In contrast, the visual cortex, a region in which AVPMNN innervation has not been observed, this phenomenon did not occur. The neuropeptide vasopressin’s involvement in this phenomenon is further demonstrated through pharmacological exposure of AVP (100nm), directly to brain ventral hippocampal slices *ex vivo*, with the strongest effect observed after 60 min of exposure, using western blot. We interpret the difference in the prompt effects of AVP *ex vivo*, compared to the delayed effects of water deprivation, to reflect the time required for water deprivation to trigger consequential increases in AVP release *in vivo*. The inhibition of AVP stimulation of GluA1 and PSD95 expression was expected to be via V1a and/or V1b receptors in ventral hippocampus. Either the V1a or the V1b antagonist ablated the AVP-stimulated increases on PSD95 or GLUA1 expression as measured by western blot (conclusion based on the absence of statistically significant differences observed, compared to control). This suggests a cooperative activation of V1a and V1b receptors is needed for the AVP-induced postsynaptic protein potentiation. However, it will require an extensive set of experiments to confirm it, which is beyond the scope of this study. However, the failure to provide a detailed explanation does not detract from the main demonstration that antagonists ablated the AVP stimulated increase in postsynaptic protein expression.

We also detected, quantitatively, an enrichment of both PSD proteins’ immunopositive puncta on locus coeruleus principal neuron dendrites, and also observed the close appositions between AVP axon terminals and both GluA1 and PSD95 puncta within tyrosine hydroxylase immunopositive dendrites, which indicates the existence of excitatory synapses established by presynaptic vasopressin containing axons. These data were obtained using the newly developed expansion microscopy method, modifying the published protocols to a lower expansion ratio which is sufficient for the purpose of this study, while facilitating handling for microscopical examination.

The synapse is a highly complex subcellular specialization involving closely apposed pre- and post-synaptic membranes and a prominent molecular machinery associated with them that enables chemical-electrical transmission between neurons, and Its dynamic adaptation in response to experience ^(27)^. In excitatory post-synapses, a set of molecules is assembled in an ultrastructural feature known as the postsynaptic density (PSD), a narrow electron – dense area subjacent to the postsynaptic membrane ^(8, 28)^. The PSD is composed of several scaffolding proteins that coordinate the trafficking and membrane insertion of α-amino-3-hidroxy-5-methyl-4-isxazolepropionate (AMPA)-or *N-*methyl-D-aspartate (NMDA)-type glutamate ionotropic receptors at the synaptic cleft, thus allowing excitatory transmitter-dependent activation of the postsynaptic neuron and stabilizing synaptic contact ^(29-35)^. This dynamic process appears to be finely tuned by various cues that include autocrine, paracrine and neurocrine signals occurring at the synaptic cleft or in the vicinity of the synapse, that can modulate molecular pathways within the PSD, allowing short-and long-term regulation of synaptic plasticity ^(36, 37)^. We chose to investigate the neurohormone vasopressin’s effect(s) on the above physiological process, through determination of changes in the expression of the glutamate AMPAR receptor subunit GluA1, a molecule which has been reported to be mainly localized within glutamatergic synapses and is hence an excitatory synapse marker ^(38)^, and PSD95, a membrane-associated guanylate kinase (MAGUK). Both PSD95 and GluA1 are crucial regulators of postsynaptic organization and function ^(39, 40)^. We focused on limbic projections domains of AVPMNN ^(9-11, 24, 41)^, in comparison with non-AVP-innervated brain areas, and under conditions in which the AVPMNN system is dramatically upregulated ^(20, 42)^. We have shown that water deprivation causes prompt enhancement of PSD95 and GluA1 levels in limbic brain areas to which AVP-glutamatergic neurons project, but not in primary visual cortex, an area in which expression of these proteins is equally abundant, but which do not receive AVPMNN innervation. Augmentation of PSD95 and GluA1 was reversible upon re-institution of water availability, indicating the state-dependency of this effect. These data reveal a novel role for AVP as a synapse organizer in limbic regions innervated by AVPMNN direct projections, a potential mechanistic link between homeostatic adversity (water deprivation) and motivational drive.

We have previously shown that water-deprived animals exposed to cat odour show diminished freezing behaviour and reduced functional output of lateral habenula, likely due to selective activation of GABAergic interneurons in this region selectively receiving glutamate-vasopressin innervation ^(11, 43)^, and suggesting that the vasopressinergic system could act as a modulator of synaptic transmission during stress responses. The present data add concrete cellular data about the possible mechanisms whereby vasopressin alters synaptic function under conditions of stress (in this case, water deprivation) by regulation of two discrete post-synaptic density proteins, PSD95 and GluA1), and extends it to other brain regions innervated by AVPMNNs, including amygdala and ventral hippocampus. This modulation could well be of physiological importance, in influencing decision-making leading to behavioural responses that facilitate survival in the face of homeostatic and environmental threats.

### Technical considerations

While the *in vivo* employment of antagonists to demonstrate specificity would be desirable, via microinjection or systematic administration, for the particular aim of this study (i.e., post-synaptic density protein expression level changes in specific brain regions), they have major limitations as systemic, or microinjection delivery cannot adequately neutralize the *in vivo* delivery of AVP directly into the synapse. This might be overcome by utilizing mice with ablation of V1a and V1b receptor genes. However, this would have to be conditional knock out as the animals might not be viable or have undergone physiological buffering (re-organization) if ablation is *ab initio*.

We continue to search for experiments to further demonstrate the mechanism of AVP-induced synaptic protein regulation. Performing similar experiments in mouse will allow genetic ablation of V1a and V1b to confirm, *in vivo*, the importance of each receptor subtype for PSD95 and GluA1 regulation by water deprivation. While this is beyond the scope of the present work, we consider it a necessary extension of the results reported here *ex vivo* (in slice preparation),and will pave the way for understanding how each receptor type signals to effect this dramatic modulation of post-synaptic protein expression.

## Acknowledgements

We thank Maurice Manning for his kind gift of arginine vasopressin, María del Carmen Cardenas-Aguayo for helping with western blot experiments. MAZ thanks UNAM-DGAPA-POSDOC fellowship. TPF and ECL thank the supported by CONACYT PhD studentship through grant CONACYT-CB-238744. RPM wishes to thank UNAM-DGAPA-PREI program for the support of his sabbatical stay at LZ lab. LZ wishes to express her gratitude to Peter Somogyi for hosting her 2nd sabbatical year (2007-2008), during which, in depth discussion on her electron microscopy observations spawned this study. This project was supported by grants: UNAM-DGAPA-PAPIIT-IN200121 & CONACYT-CB-238744 and 283279 (LZ); CONACYT A1-S-8731 (LIE); NIMH-IRP, NIH, ZIAMH0002386 (LEE).

## Supplementary information

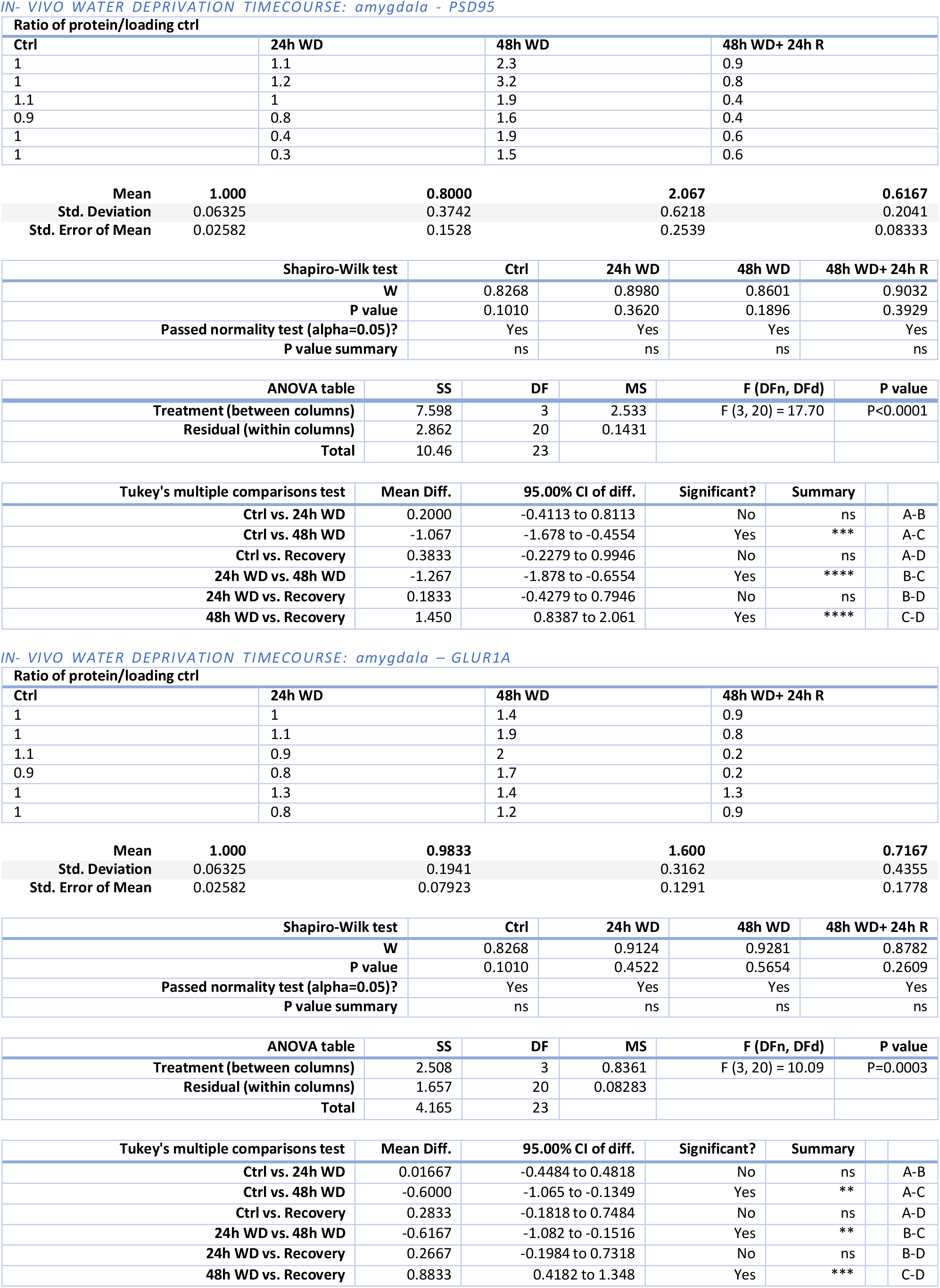
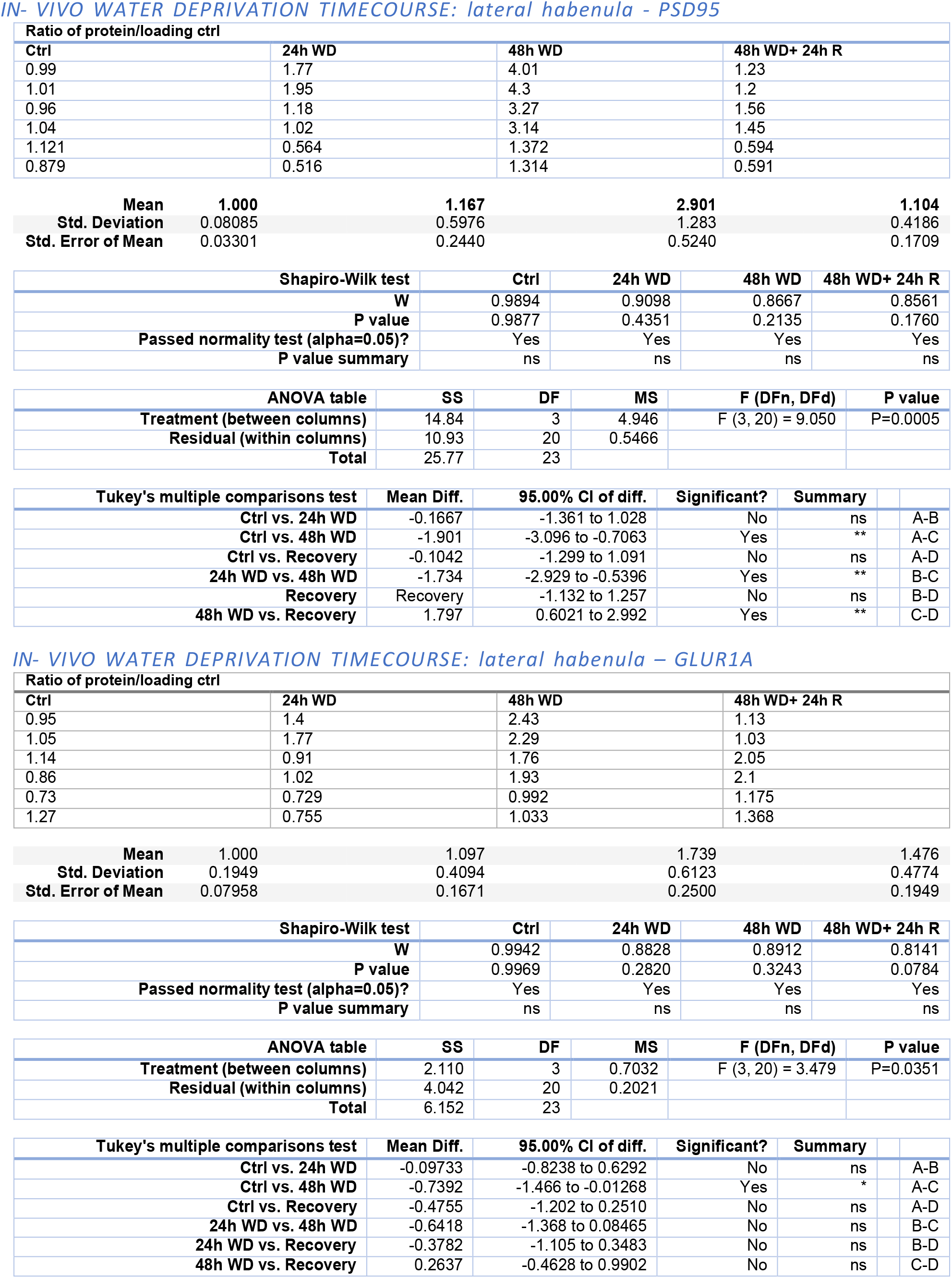
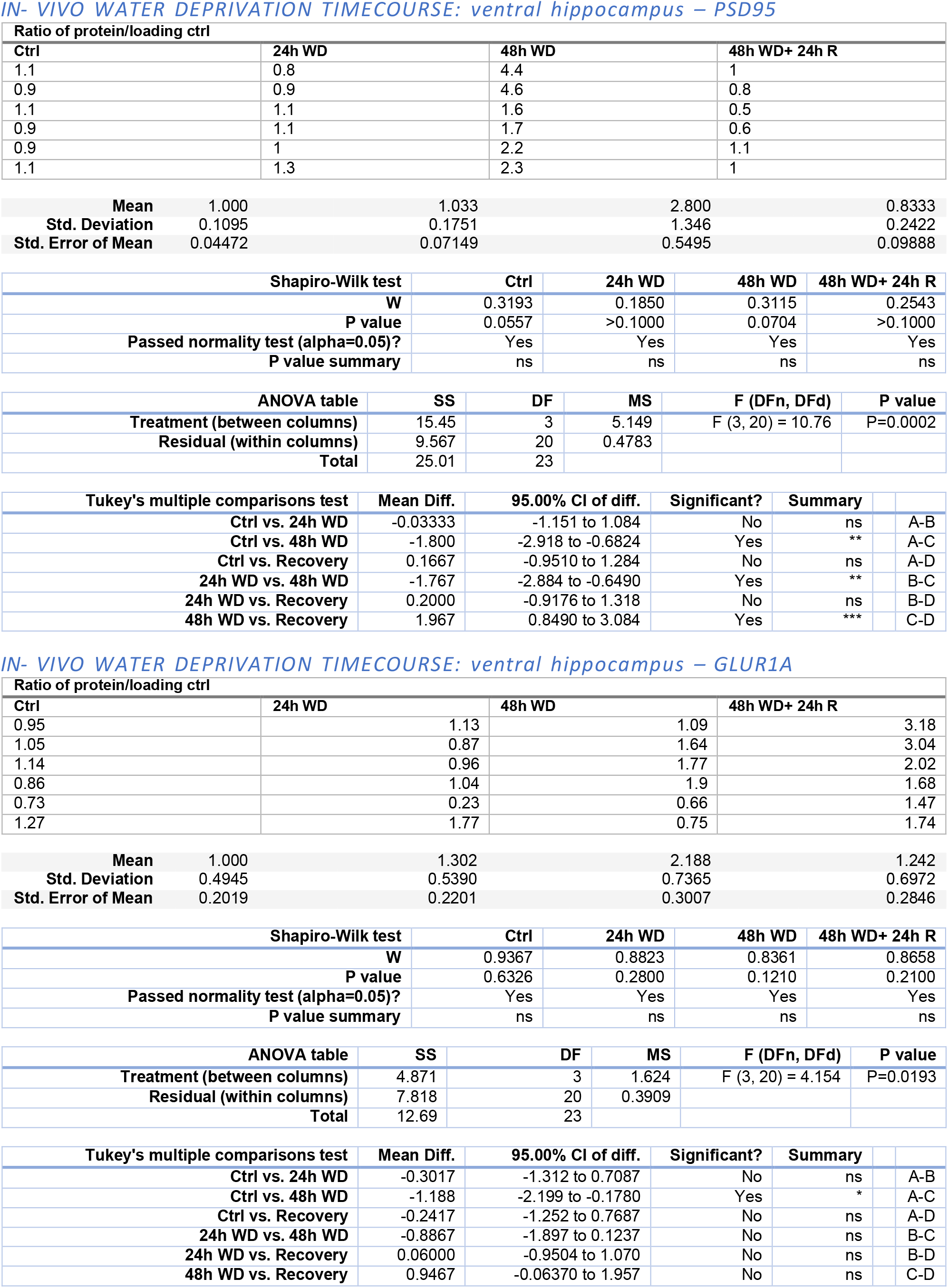
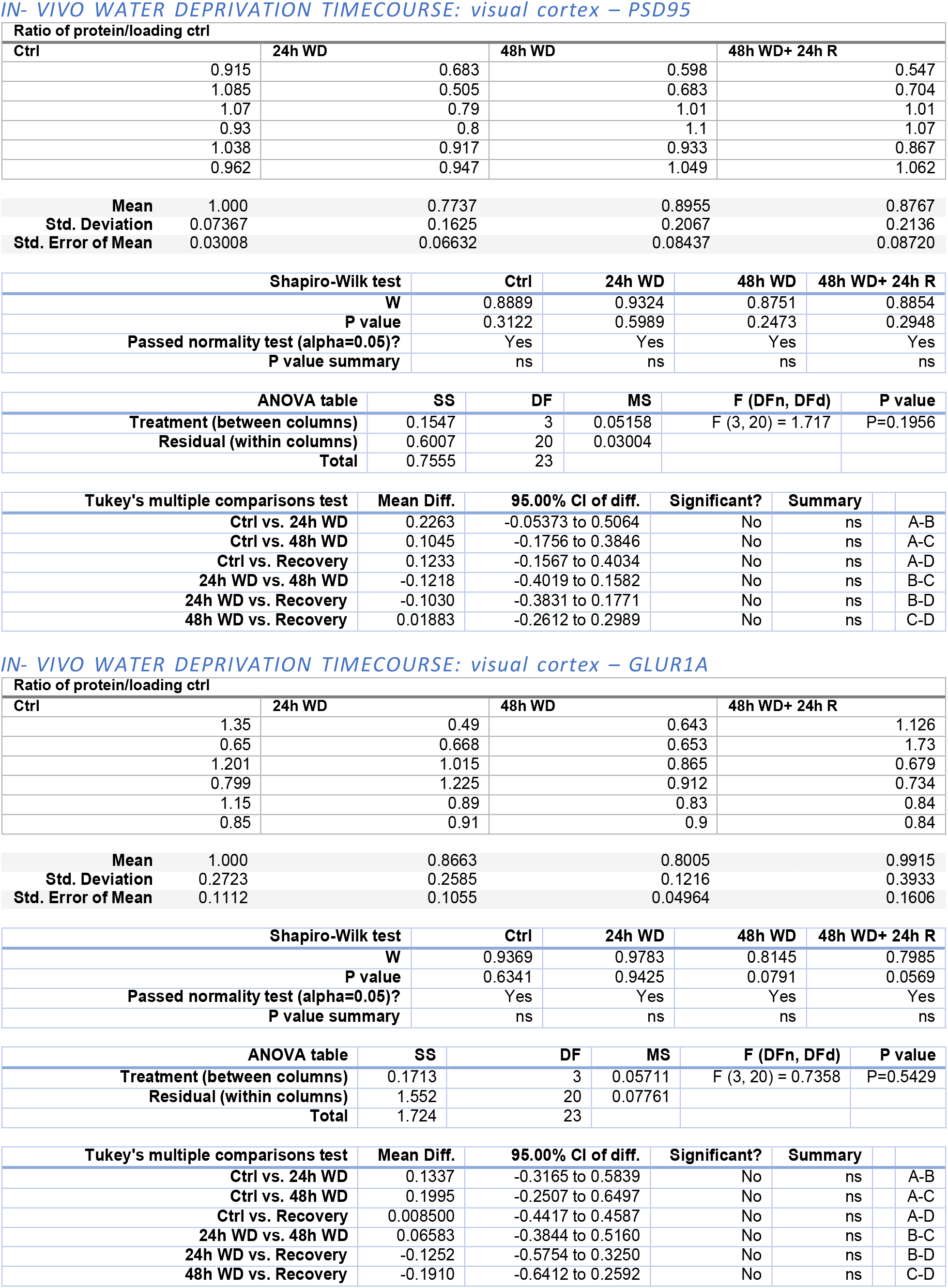
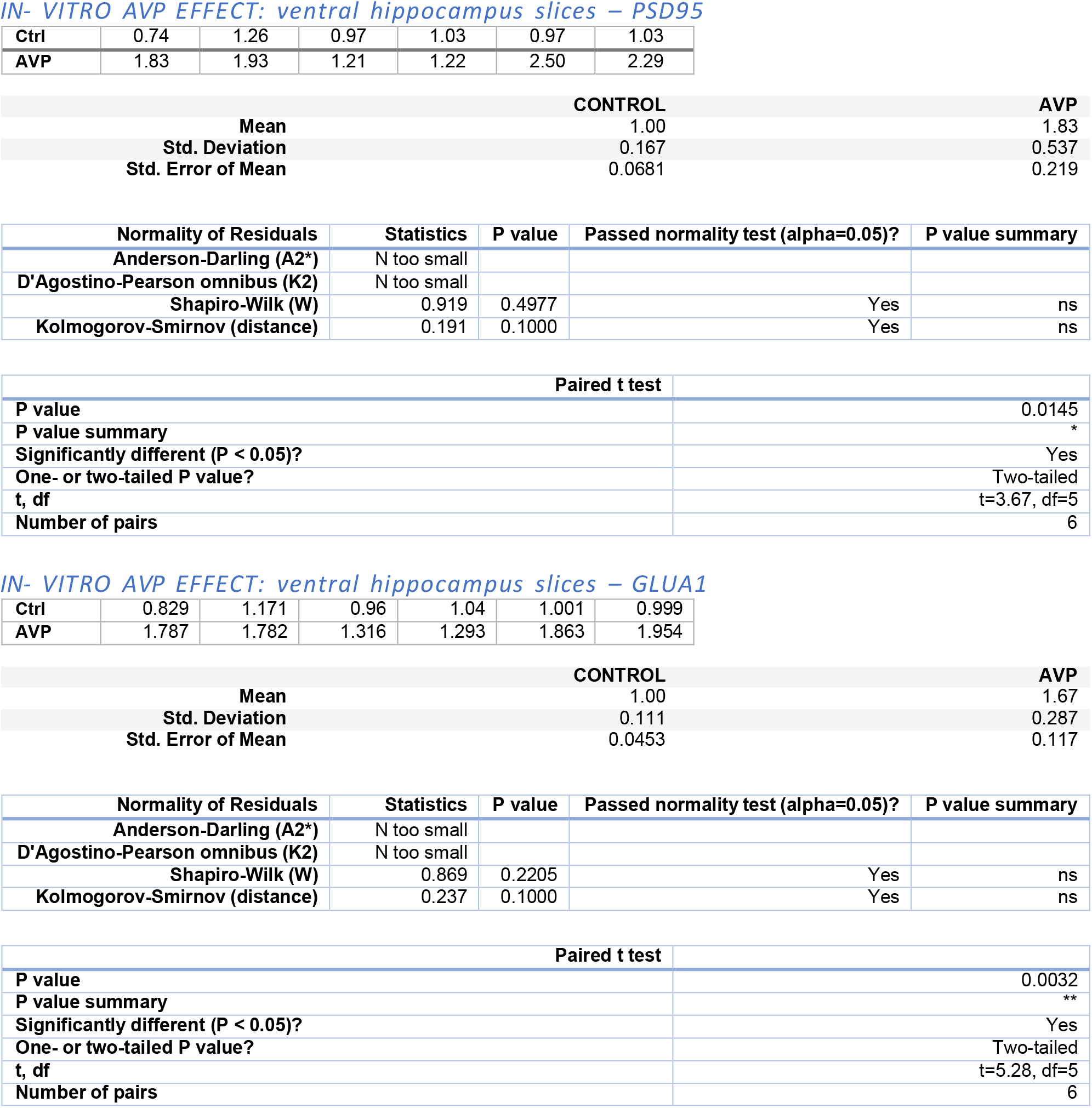
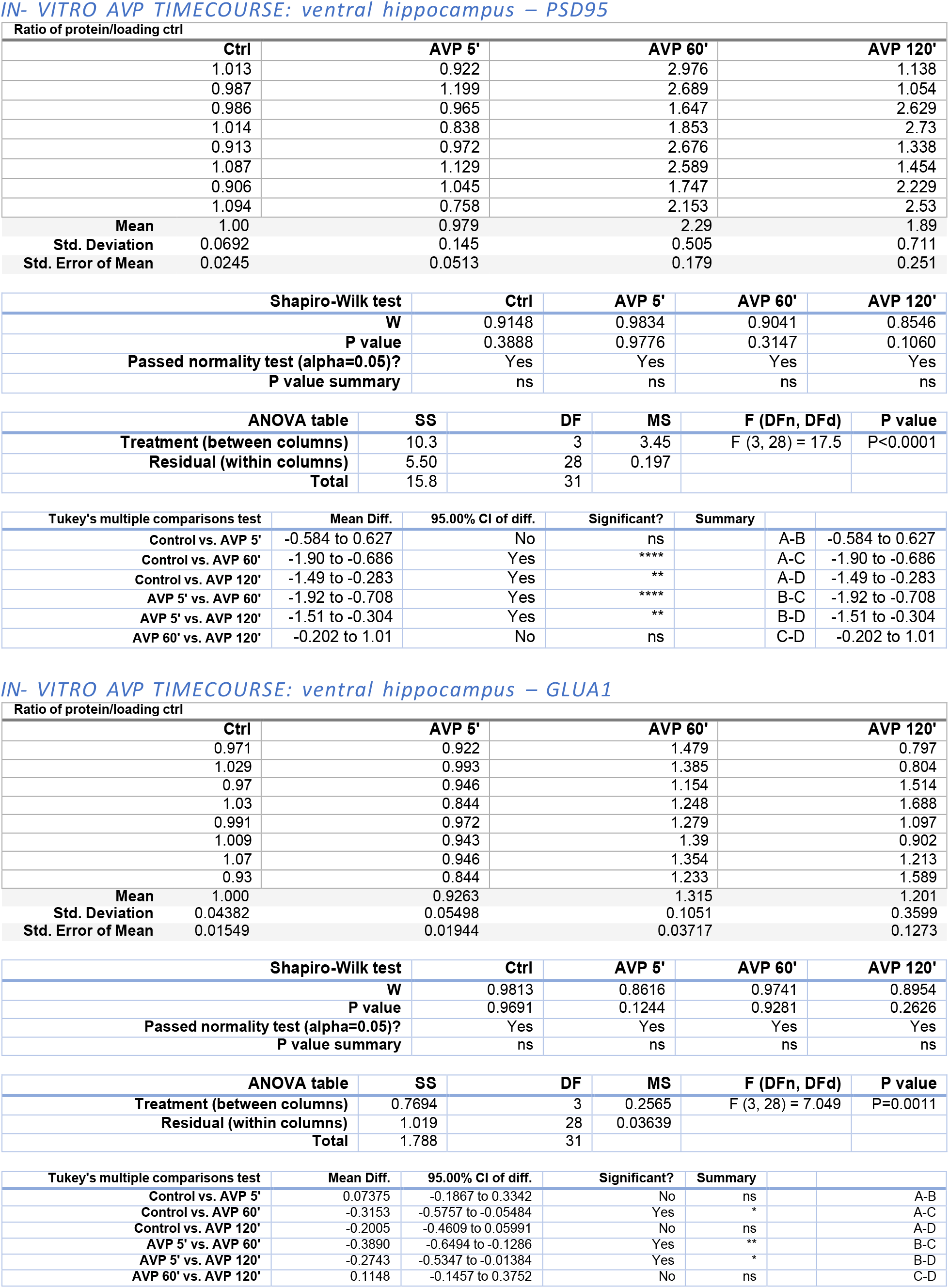
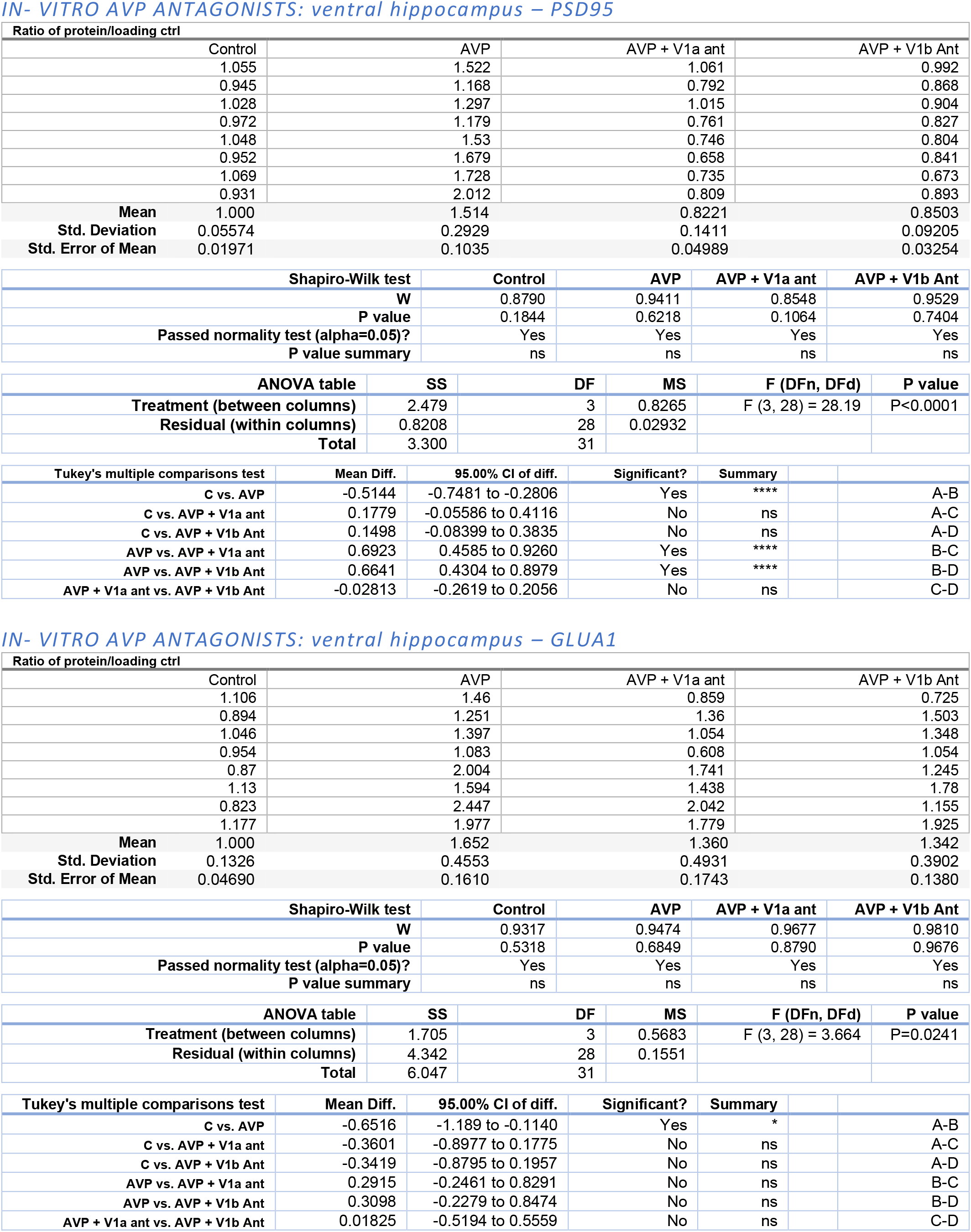
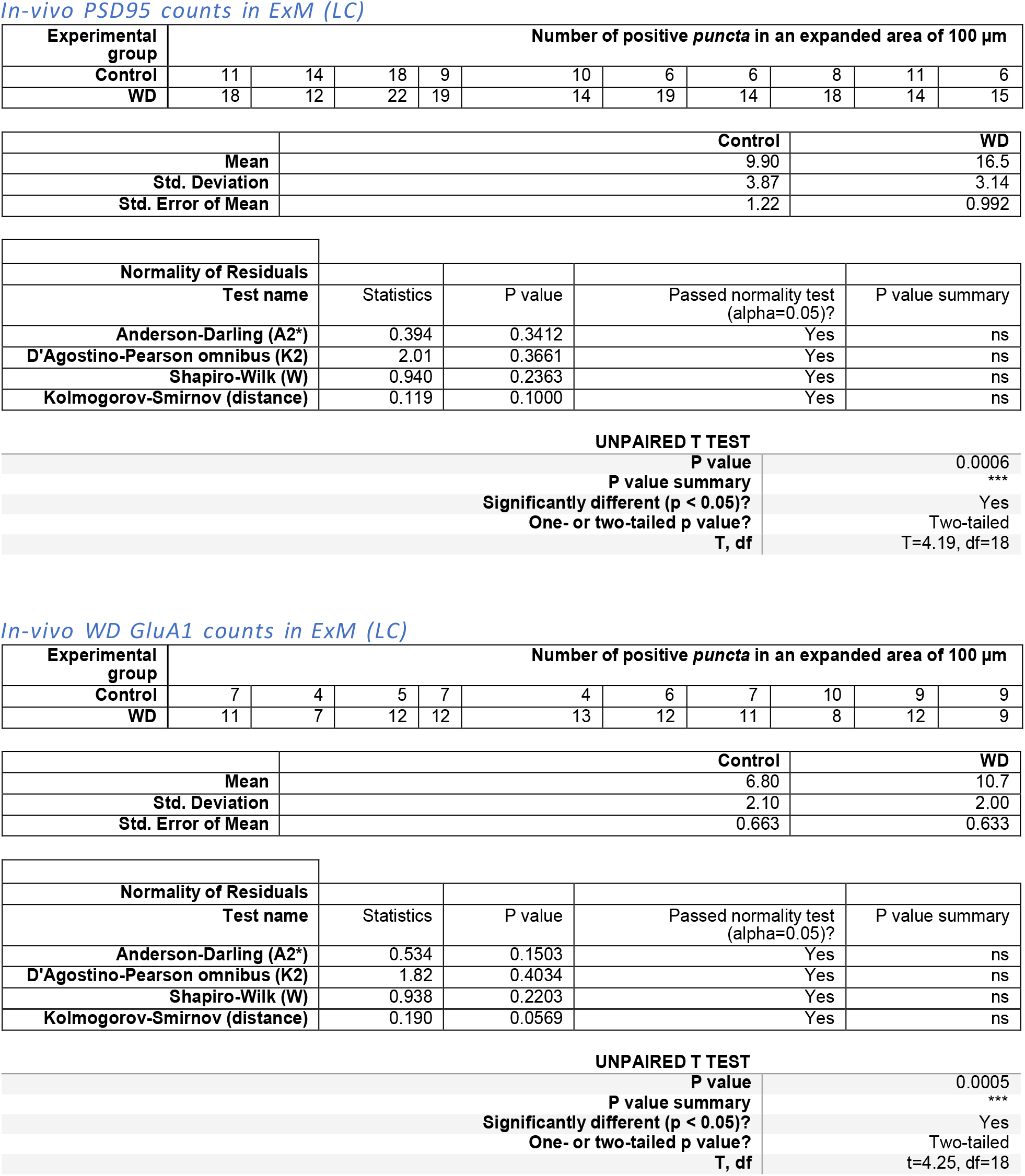

